# A curated collection of *Klebsiella* metabolic models reveals variable substrate usage and gene essentiality

**DOI:** 10.1101/2021.10.10.463858

**Authors:** Jane Hawkey, Ben Vezina, Jonathan M. Monk, Louise M. Judd, Taylor Harshegyi, Sebastián López-Fernández, Carla Rodrigues, Sylvain Brisse, Kathryn E. Holt, Kelly L. Wyres

## Abstract

The *Klebsiella pneumoniae* species complex (KpSC) is a set of seven *Klebsiella* taxa which are found in a variety of niches, and are an important cause of opportunistic healthcare-associated infections in humans. Due to increasing rates of multi-drug resistance within the KpSC, there is a growing interest in better understanding the biology and metabolism of these organisms to inform novel control strategies. We collated 37 sequenced KpSC isolates isolated from a variety of niches, representing all seven taxa. We generated strain-specific genome scale metabolic models (GEMs) for all 37 isolates and simulated growth phenotypes on 511 distinct carbon, nitrogen, sulphur and phosphorus substrates. Models were curated and their accuracy assessed using matched phenotypic growth data for 94 substrates (median accuracy of 96%). We explored species-specific growth capabilities and examined the impact of all possible single gene deletions on growth in 145 core carbon substrates. These analyses revealed multiple strain-specific differences, within and between species and highlight the importance of selecting a diverse range of strains when exploring KpSC metabolism. This diverse set of highly accurate GEMs could be used to inform novel drug design, enhance genomic analyses, and identify novel virulence and resistance determinants. We envisage that these 37 curated strain-specific GEMs, covering all seven taxa of the KpSC, provide a valuable resource to the *Klebsiella* research community.

## Introduction

*Klebsiella pneumoniae* is a ubiquitous bacterium that inhabits a variety of host- and non-host associated environments and is a major cause of human disease. It is an opportunistic pathogen and a significant contributor to the spread of antimicrobial resistance globally (Pendleton et al. 2014; Navon-Venezia et al. 2017; Thorpe et al. 2021). Multi-drug resistant *K. pneumoniae* with resistance to the carbapenems (the ‘drugs of last resort’) cause infections that are extremely difficult to treat and are considered an urgent public health threat (Pendleton et al. 2014). Understanding the biology and ecological behaviour of these organisms is essential to inform novel control strategies.

The past 6-7 years have seen an explosion of *K. pneumoniae* comparative genomics studies, revealing numerous insights into its epidemiology, evolution, pathogenicity and drug-resistance, and informing a genomic framework that facilitates surveillance and knowledge generation (recently summarised in (Wyres et al. 2020)). It is now clear that isolates identified as *K. pneumoniae* through standard microbiological identification techniques actually comprise seven distinct closely related taxa known as the *K. pneumoniae* species complex (KpSC): *K. pneumoniae sensu stricto, Klebsiella variicola* subsp. *variicola, K. variicola* subsp. *tropica, Klebsiella quasipneumoniae* subsp. *quasipneumoniae, K. quasipneumoniae* subsp. *similipneumoniae, Klebsiella quasivariicola* and *Klebsiella africana* (Gorrie et al. 2017; Long et al. 2017; Rodrigues et al. 2019; Wyres et al. 2020). *K. pneumoniae sensu stricto* accounts for the majority of human infections and is therefore the most well-studied of these organisms.

Each individual *K. pneumoniae* genome encodes between 5000 and 5500 genes; ∼2000 are conserved among all members of the species (core genes) and the remainder vary between individuals (accessory genes) (Holt et al. 2015). The total sum of all core and accessory genes is estimated to exceed 100,000 protein coding sequences that can be assigned to various functional categories, many of which are not well-characterised. For example, the diversity, mechanism and phenotypic impact of antimicrobial resistance genes, accounting for 1% of the total gene pool, is well understood. In contrast the functional implications of metabolic genes, which account for the largest single fraction of the gene-pool (37%) (Holt et al. 2015), are relatively poorly understood. The sheer number of genes in this category suggests that substantial metabolic variability exists within the KpSC, a hypothesis supported by two studies that have generated growth phenotypes for multiple isolates (Brisse et al. 2009; Blin et al. 2017). However, these data are limited by the number and variety of substrates tested and it is difficult to consolidate the genotype data in the context of these phenotypes. Moreover, these phenotyping methods are slow, expensive, and non-scalable across large numbers of isolates.

Genome-scale metabolic modelling represents a powerful approach to bridge the gap between genotypes and phenotypes. Drawing on the accumulated biochemical knowledge-base, it is possible to infer the metabolic network of an individual organism from its genome sequence and subsequently apply *in silico* modelling approaches to predict its metabolic capabilities (growth phenotypes) (O’Brien et al. 2015). Such models allow exploration of metabolic diversity (Monk et al. 2013; Seif et al. 2018; Bosi et al. 2016), prediction the impact of gene deletions or the response to drug exposure (Tong et al. 2020), identification of novel virulence factors or drug targets (Ramos et al. 2018; Bartell et al. 2017; Zhu et al. 2018), and optimisation for the production of industrially-relevant compounds. (Li et al. 2016; Jung et al. 2015).

To-date, two curated and validated single strain genome-scale metabolic models (GEMs) have been reported for *K. pneumoniae*. The first was generated for the MGH78578 laboratory strain and published in 2011 (model ID iYL1228) (Liao et al. 2011). It comprised 1228 genes, 1188 enzymes and 1970 reactions, and was validated by comparison of *in silico* growth predictions to true phenotypes generated for 171 substrates using a Biolog phenotyping array. The estimated accuracy of iYL1228 was 84% when comparing to Biolog growth phenotypes. A second *K. pneumoniae* GEM, for laboratory strain KPPR1, was published in 2017 (model ID iKp1289) (Henry et al. 2017). This model contained 1289 genes and 2145 reactions. The KPPR1 model was found to be 79% accurate when compared to Biolog phenotype data in terms of predicting substrate-growth phenotypes. More recently, Norsigian and colleagues (Norsigian et al. 2019a) reported non-validated draft GEMs for 22 antimicrobial-resistant *K. pneumoniae* clinical isolates built from the iYL1228 model via a subtractive approach. Subsequent *in silico* growth predictions indicated variability between isolates in terms of carbon, nitrogen and sulfur but not phosphorus utilisation. There was evidence that nitrogen substrate usage could be used to classify strains associated with distinct drug-resistance phenotypes. However, none of these models were experimentally validated.

Here, we present an updated version of the MGH78578 GEM in addition to novel GEMs for 36 KpSC strains, including representatives of all seven taxa in the species complex. We curate and validate the models using a combination of Biolog growth assays and additional targeted growth phenotype data, resulting in a median accuracy of 96%. We define the core reactomes of *K. pneumoniae* and the broader species complex, and identify species-specific metabolic capabilities. We then explore these models to identify strain-specific gene essentiality and metabolic pathway redundancy across growth on 145 core carbon substrates.

## Results

### Completed KpSC Genomes

We collated 37 previously described isolates from the KpSC complex, including at least one representative per taxon (Blin et al. 2017; Rodrigues et al. 2019). The collection spanned a variety of sequence types (STs) within species with more than one strain, and represented a wide range of isolation sources (including human host-associated, water and the environment). The strains were geographically and temporally diverse, sampled from five continents and with isolation dates spanning 1935 - 2010 (**Supplemental Table 1**).

Eight strains had previously-published complete genome sequences available, and we generated complete genome sequences for the remaining 29 strains using a combination of short- and long-read sequencing (see **Methods**). The median genome size was 5.5 Mbp (range 5.1 - 6.0 Mbp) with a median of 5145 genes (range 4798 - 5704 genes). The majority of strains carried at least one plasmid (n=29, 78%), with seven strains carrying five or more plasmids.

### Model generation, curation and validation

Using these completed genomes we created strain-specific GEMs, initially using the curated MGH78578 GEM (iYL1288) as a reference to identify conserved genes and reactions, followed by manual curation (see **Methods**). The latter was enabled by the availability of matched phenotype data (Blin et al. 2017) indicating the ability of each strain to grow in minimal media supplemented with each of 94 distinct sole carbon substrates for which we were able to predict growth in silico using the GEMs (**Supplemental Table 2**). Our phenotypic data included 12 carbon substrates for which growth was demonstrated for at least one strain and for which the corresponding metabolite transport and/or processing reactions were not present in the original iYL1288 model. Literature searches were undertaken to identify the putatively responsible candidate genes and reactions for GEM inclusion. For example, all strains were able to utilise palatinose as a carbon substrate; the reaction required to catabolise this compound was added based on the presence of core genes with ≥99% nucleotide homology to *aglAB* (that encode AglAB), which has been shown to catabolise palatinose in *K. pneumoniae* (Thompson et al. 2001) (**Supplemental Table 3**). When the model-based predictions and our phenotypic growth data disagreed, we attempted to correct the models by identifying alternative pathways from the literature or homologous genes in other *Klebsiella* or Enterobacteriaceae species with sufficient evidence to allow inclusion in our models (see **Methods, Supplemental Table 3**). Overall, we added 49 genes and 56 reactions across all models.

The final curated, validated models were highly accurate for the prediction of growth phenotypes (median accuracy 95.7%, range 88.3 - 96.8%, **Supplemental Table 1**). The majority (87%) of the discrepancies were false positives, where the model predicted growth on a carbon substrate but we did not observe any phenotypic growth. False positives usually occur due to gene regulation, where strains carry the genes encoding the enzymes required to import and metabolise a substrate, however these genes are not expressed during the phenotypic growth experiments. False positives can also be related to technical issues with measuring metabolic phenotypes, e.g. the LOD, sensitivity of growth detection, and use of correct standards for measurements (Ibarra et al. 2002). Every model had at least one false positive (median 4, range 1 – 11, **Supplemental Table 1**) across 31 different carbon substrates. The most common false positive calls were predicted growth in 2-oxoglutarate (n=35 strains), ethanolamine (n=29), L-ascorbate (n=28) and 3-hydroxycinnamic acid (n=20); false positive calls for the remaining 27 carbon substrates were associated with ≤6 strains each (**Supplemental Table 4**).

Five carbon substrates had at least one strain with a false negative call, where the model did not predict growth but we observed a growth phenotype: L-tartaric acid (n=12 strains), L-lyxose (n=5), L-sorbose (n=2), propionic acid (n=2) and L-galactonic acid-gamma-lactone (n=1) (**Supplemental Table 4**). In such cases it is assumed that the models are missing information required to optimise for growth on these substrates (Orth et al. 2012). Despite thorough literature and database searches, we were unable to identify alternate biological pathways that could plausibly fill these gaps in the models. This was particularly notable among the five *K. quasipneumoniae* subsp. *quasipneumoniae* strains, which all had false negative predictions for L-lyxose utilisation. These genomes were each missing *sgaU* (KPN_04590), which was present in all other KpSC genomes and encodes an enzyme that converts L-ribulose-5-phosphate to L-xylulose-5-phosphate. We were unable to detect any other proteins belonging to this enzyme class or carrying similar domains. As the phenotypic results indicated that all *K. quasipneumoniae* subsp. *quasipneumoniae* can utilise L-lyxose, we hypothesise that they must contain unknown functional orthologue/s to *sgaU*, which can perform isomerase activity on L-ribulose 5-phosphate.

### Novel GEMs reveal species- and strain-specific metabolic diversity

Our strain collection provided us with a novel opportunity to compare predicted metabolic functionality between all seven taxa within the KpSC. Overall there were median 1219 genes and 2294 reactions in each curated strain-specific GEM (range 1190 - 1243 and 2283 - 2305 respectively), representing median 23.6% of all coding sequences in each genome (**Supplemental Table 1**). Each species had ∼1200 core model genes and ∼2200 core reactions (**Table 1**), with a slight decreasing trend with increasing sample size. Conversely, the total number of distinct reactions detected among the best represented species, *K. pneumoniae* (2312, n=20 genomes) was higher than those detected among each of the species represented by fewer genomes (2299 in *K. quasipneumoniae* subsp. *quasipneumoniae*; 2307 in both *K. quasipneumoniae* subsp. *similipneumoniae* and *K. variicola* subsp. *variicola*). In terms of the reactions themselves, the vast majority were core across all species (**Fig. 1**), however there was notable variability in reactions associated with carbohydrate metabolism, for which 16% (n=37/234) were not conserved across all models (**Fig. 1**). Among these variable reactions we identified three involved in the N-acetylneuraminate pathway (ACNAMt2pp, ACNML and AMANK) which were species-specific and were found to be core in *K. quasipneumoniae* subsp. *similipneumoniae* while absent from all other genomes.

**Table 1:** Summary of genomes and the core elements of the GEMs.

| Species | # genomes | # STs | # model genes (core) | # reactions (core) | # phenotypes (core) |
| --- | --- | --- | --- | --- | --- |
| *K. pneumoniae* | 20 | 18 | 1202 - 1243 (1183) | 2288 - 2305 (2276) | 277 - 282 (277) |
| *K. quasipneumoniae* subsp. *quasipneumoniae* | 5 | 5 | 1197 - 1209 (1190) | 2283 - 2289 (2283) | 270 - 274 (268) |
| *K. quasipneumoniae* subsp. *similipneumoniae* | 5 | 5 | 1200 - 1220 (1194) | 2283 - 2299 (2287) | 273 - 280 (273) |
| *K. variicola* subsp. *variicola* | 4 | 4 | 1212 - 1227 (1214) | 2294 - 2301 (2299) | 279 - 282 (279) |
| *K. africana* | 1 | 1 | 1216 | 2289 | 279 |
| *K. quasivariicola* | 1 | 1 | 1228 | 2299 | 279 |
| *K. variicola* subsp. *tropica* | 1 | 1 | 1237 | 2310 | 281 |

**Figure 1:**
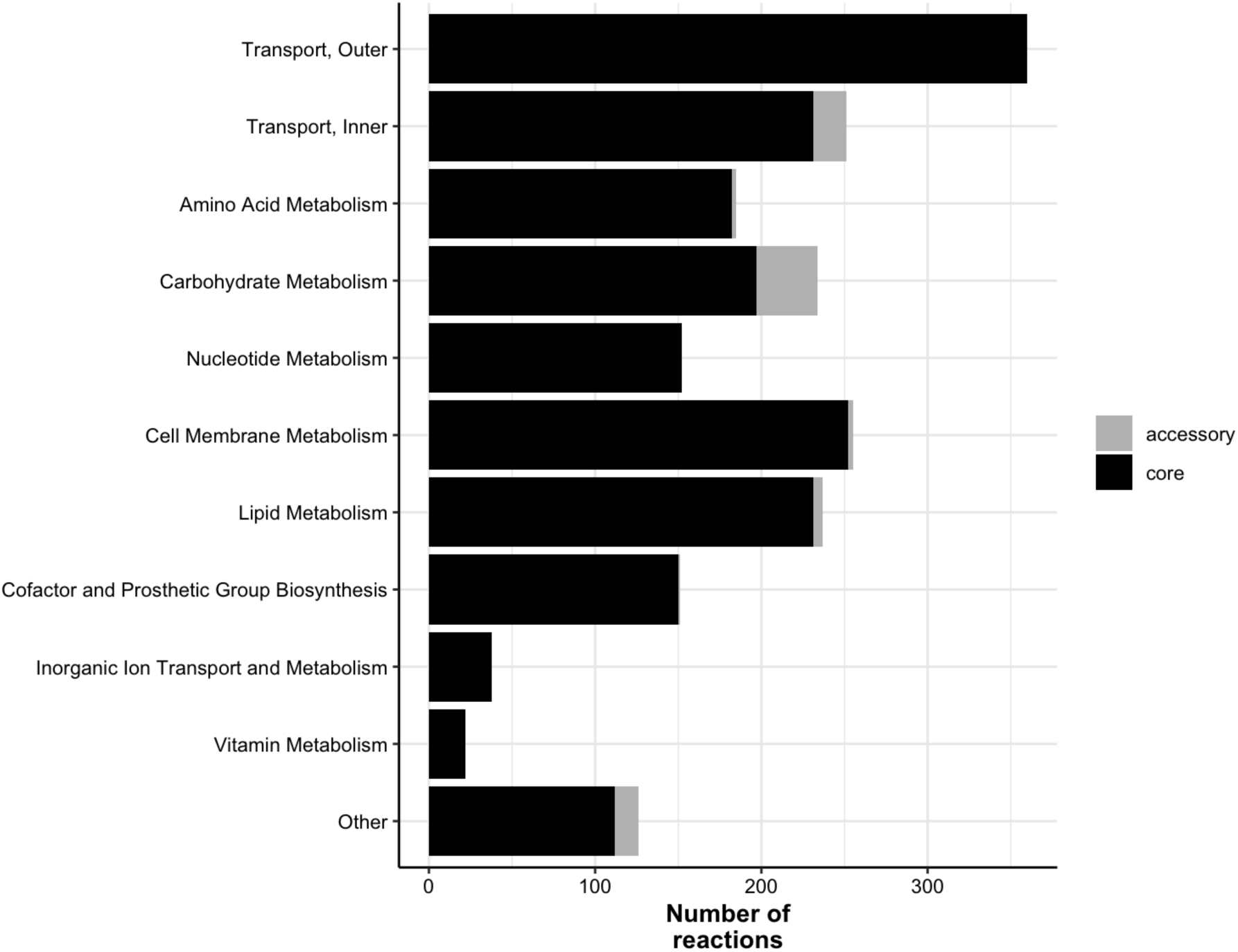
Number of model reactions by category. Bars are coloured to indicate core reactions (black, conserved in all strains) and accessory reactions (grey, variably present).

We simulated growth on 511 substrates as the sole sources of either carbon (n=272), nitrogen (n=155), phosphorus (n=59) or sulfur (n=25) (see **Methods, Supplemental Table 2**). A total of 224 (44%) were unable to support growth for any strain (carbon=107, nitrogen=87, phosphorus=15, sulfur=15). Overall the number of core growth-supporting phenotypes was very similar across taxa, with a median of 279 (range 268 - 281, **Table 1**). Of the 287 that were predicted to support growth for at least one strain, 262 were conserved across all 37 strains (carbon=145, nitrogen=64, phosphorus=43, sulfur=10), with only 25 (5%) substrates variable between strains. Substrates that could be utilised as a carbon source had the most variation, with 7% of carbon substrates displaying variable predicted growth phenotypes by strain (**Fig. 2**). This was in stark contrast to substrates used as a source of sulfur, where no variation was observed (**Fig. 2**).

**Figure 2:**
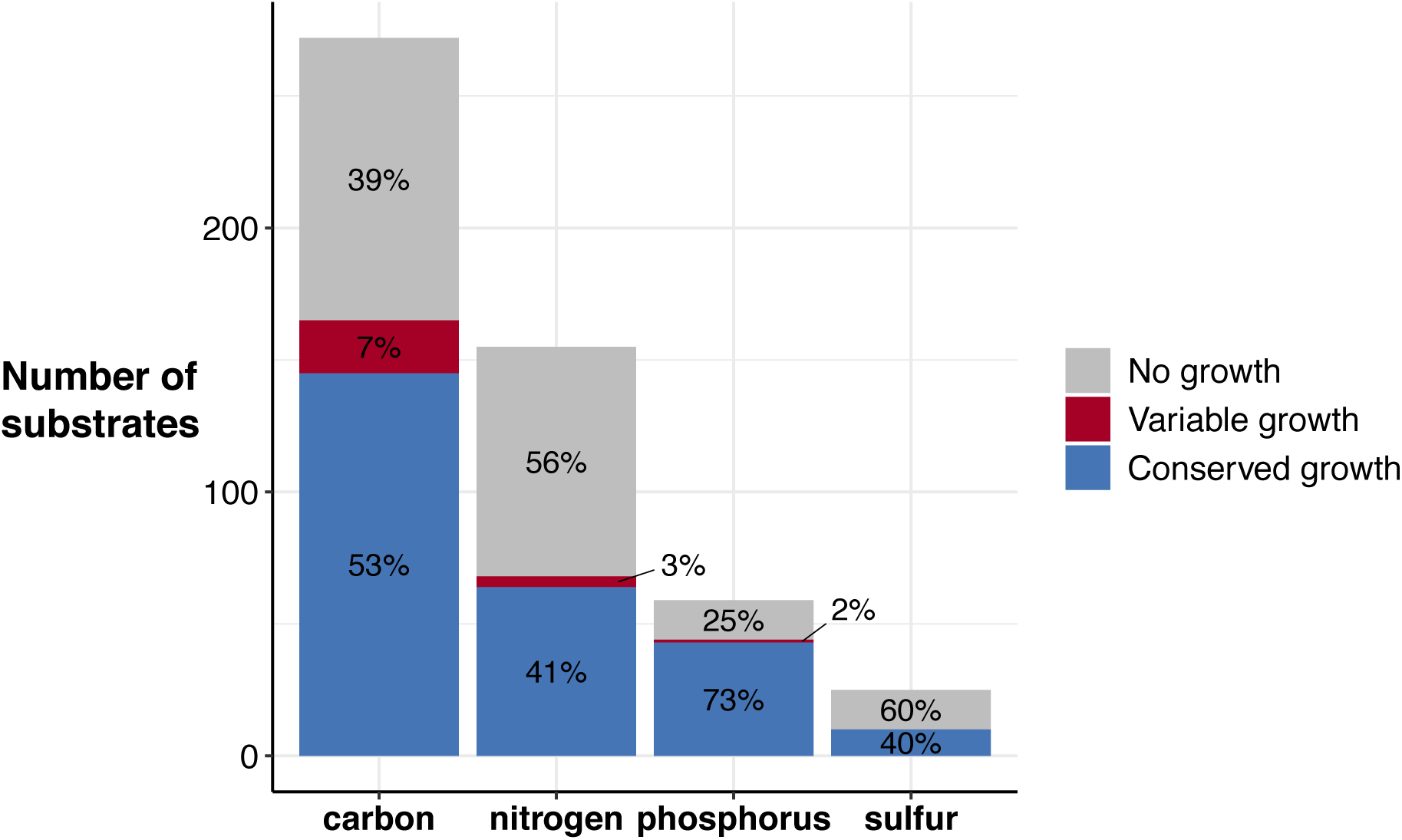
Predicted substrate utilisation by type. Bar height indicates number of substrates for each type, with segments coloured to indicate those associated with no growth for any strain (grey), variable growth (red) and conserved growth (blue). Percentages are indicated within each segment.

Amongst the 20 variable carbon substrates, there was some species-specific variation. Six of these reflect core growth capabilities in all but one of the seven species (3-hydroxycinnamic acid, 3-(3-hydroxy-phenyl)propionate, D-arabitol, L-ascorbate, L-lyxose, tricarballylate, **Fig. 3**). In the case of tricarballylate, we identified a new pathway which was absent from the original *K. pneumoniae* MGH78578 model: all KpSC species except for *K. pneumoniae* carried the *tcuABC* operon, which encodes the enzymes responsible for oxidising tricarballylate to cis-aconitate (Lewis et al. 2009) via the TCBO reaction (**Fig. 3**). In contrast, all KpSC were able to utilise L-ascorbate with the exception of *K. quasipneumoniae* subsp. *quasipneumoniae*, where all five genomes were lacking the *ulaABC* operon encoding the transport reaction ASCBptspp (**Fig. 3**). This reaction converts L-ascorbate into L-ascorbate-6-phosphate as it is transported into the cytosol (Zhang et al. 2003).

**Figure 3:**
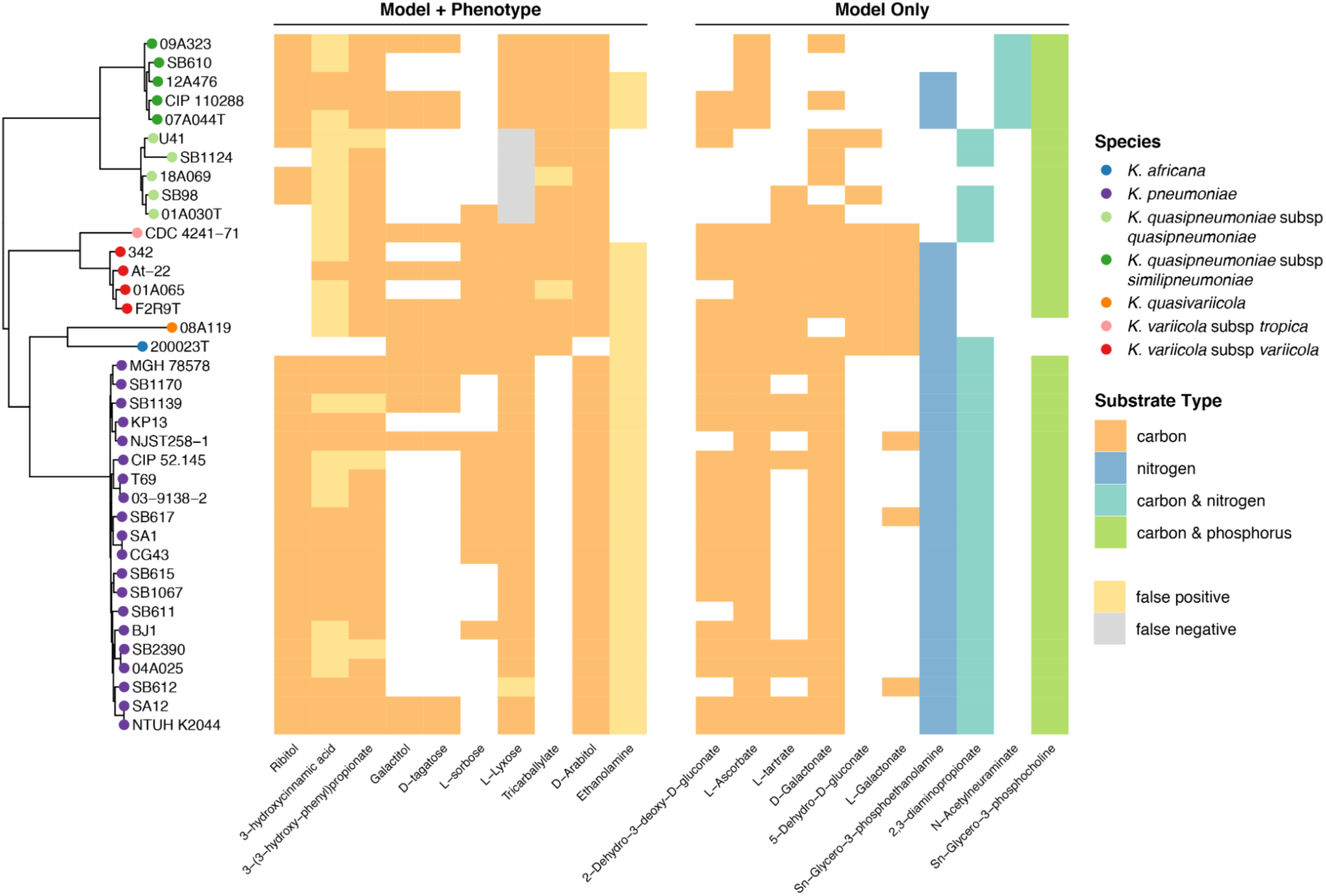
Variable growth phenotypes across all seven taxa in KpSC. Left, core gene phylogeny for all 37 strains, with tips coloured by species as per legend. Middle, heatmap of variable substrates for which both phenotypic growth results and model predicted results were available. White indicates no growth, colour indicates growth. False positive calls are shown in yellow, and false negative calls in grey (as per legend). Right, heatmap of variable substrates for which only model predictions were available. White indicates no growth, colour indicates growth, with substrate type indicated as per legend.

The remaining 14 variable carbon substrates were specific to five or fewer strains. For example, sn-glycero-3-phosphocholine could be utilised by all strains as a carbon and phosphorus substrate, except for the single *K. africana* and *K. quasivariicola* representatives, which share a common ancestor in the core-gene phylogenetic tree (**Fig. 3**). Both of these genomes lacked *ugpQ*, encoding the enzymes required to convert sn-glycero-3-phosphocholine into sn-glycero-3-phosphate and ethanolamine (Brzoska and Boos 1988). There was only a single carbon substrate, N-acetylneuraminate, which supported growth for all *K. quasipneumoniae* subsp. *similipneumoniae*, due to the presence of the *nan* operon (Vimr and Troy 1985), encoding the proteins required to catalyse the ACNAMt2pp, ACNML and AMANK reactions, which were absent in all the other species (**Fig. 3**).

### Single gene knockout simulations reveal variable gene essentiality

Strain-specific GEMs provide an unparalleled opportunity to simulate the impact of single gene knockout mutations for diverse strains. As carbon substrates were associated with the greatest amount of variation, we focused on the impact of single gene knockouts in this group. For each strain we simulated the impact of deletion of each unique gene in its GEM on growth in each of the core carbon substrates (those predicted to support growth of all strains, n=145), resulting in 6,544,865 unique simulations (**Supplemental Table 5**). Among these simulations, 639,365 (9.8%) were predicted to result in a loss of growth phenotype.

In order to compare the diversity of knock-out phenotypes between strains, we focused on simulations representing core gene-substrate combinations (n=164,285 gene-substrate combinations; 1133 genes that were present in all GEMs x 145 substrates) and excluded those representing non-core gene-substrate combinations (n=19,140 combinations), because the former can be directly compared for all strains whereas the latter cannot (by definition not all strains harbour all of the genes). A total of 146,385 core gene-substrate combinations (89.1%) resulted in no loss of growth phenotype in any strain, while 7170 (10.5%) combinations resulted in a loss of growth phenotype in all strains. At the gene level, 807 genes (71.2%) were not predicted to be essential for growth for any substrate in any strain, and just 57 genes (5.0%) were predicted to be essential for all substrates in all strains. The latter were associated with 194 distinct reactions (1-32 reactions each, median=1, **Supplemental Table 6**), encompassing 8 subsystem categories: cell membrane metabolism (n=76 reactions), lipid metabolism (n=42), amino acid metabolism (n=33), transport, inner- (n=29) or outer-transport (n=6), nucleotide metabolism (n=5), carbohydrate metabolism (n=2), and cofactor and prosthetic group biosynthesis (n=1).

Gene essentiality varied considerably by strain, with reasonable consistency within species. The number of core gene-substrate combinations predicted to result in a loss of growth phenotype ranged from 0 to 519 (median=143, **Fig. 4**) and the number of core genes resulting in a phenotype on at least one growth substrate ranged from 0 to 15 (median=3). The vast majority of these genes (31 of 36 unique genes, 86.1%) were associated with loss of growth phenotypes for ≤6 substrates, with minimal variation in the total number of substrates among those strains that were impacted. In contrast, a small number of genes were associated with loss of growth for all or almost all substrates for some strains (4 genes, 11.1%, each impacting ≥143 substrates per strain, **Fig. 4**).

**Figure 4:**
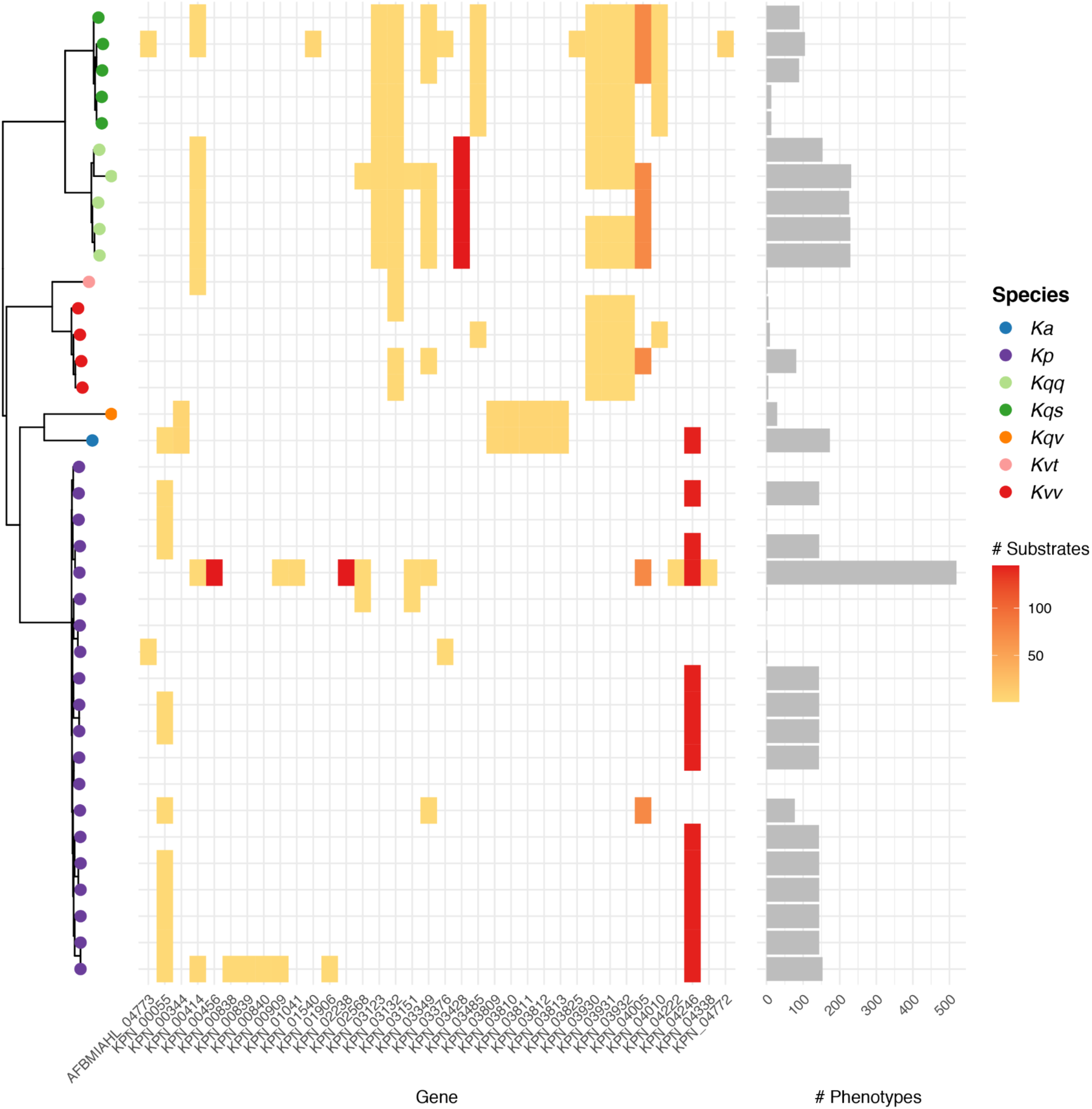
Variable loss of growth phenotypes. Left, core gene phylogeny as per **Fig 3**, with tips coloured by species as indicated in legend: Ka, *K. africana*; Kp, *K. pneumoniae*; Kqq, *K. quasipneumoniae* subsp. *quasipneumoniae*; Kqs, *K. quasipneumoniae* subsp. *similipneumoniae*; Kqv, *K. quasivariicola;* Kvt, *K. variicola* subsp. *tropica*; Kvv, *K. variicola* subsp. *variicola*. Middle, heatmap showing core genes for which variable loss of growth phenotypes were predicted (columns). Shading indicates the number of substrates where loss of growth was predicted for each strain (rows) as per the scale legend. Right, bars show the total number of loss of growth phenotypes predicted for each strain.

We further investigated the core gene deletions predicted to result in loss of growth phenotypes for ≥143 substrates in only a subset of strains, beginning with an apparent *K. quasipneumoniae* subsp. *quasipneumoniae* species-specific phenotype. The associated gene, KPN_03428, encodes the enzyme for catalysis of two reactions in the models: CYSDS (cysteine desulfhydrase) and CYSTL (cystathionine b-lyase), the latter of which may also be encoded by KPN_01511 (*malY*). Notably, *malY* was present in all other models but absent from all *K. quasipneumoniae* subsp. *quasipneumoniae* (closest bi-directional BLASTp hit had 30.07% identity, well below the threshold required for inclusion as a homolog and considerably lower than the expected divergence between KpSC *s*pecies (Holt et al. 2015)), and no alternate genes encoding putative cystathionine b-lyases could be identified by search of the KEGG database, indicating a lack of genetic redundancy for these reactions. Direct comparison of the *K. quasipneumoniae* subsp. *quasipneumoniae* 01A030T chromosome to *K. pneumoniae* MGH78578 revealed that the former harboured a ∼5 kbp deletion relative to the latter, spanning the *zntB, malY* and *malX* genes as well as part of *malI*. The lack of *malY* (KPN_01511) in combination with the KPN_03428 deletion resulted in predicted loss of ability to produce three key metabolites (L-homocysteine, ammonium and pyruvate) and ultimately the predicted loss of biomass production. This deletion was replicated in all five *K. quasipneumoniae* subsp. *quasipneumoniae* strains. Inspection of an additional 149 publicly available *K. quasipneumoniae* subsp. *quasipneumoniae* genome assemblies (see **Methods**) found this region to be present in only 37 genomes (24%), suggesting that the most recent common ancestor of this species is lacking this region, with occasional re-acquisition in some lineages.

Unlike the KPN_03428 deletion, deletion of KPN_04246 resulted in predicted loss of growth phenotypes for all 145 substrates for the single *K. africana* strain plus 13 of 20 K. pneumoniae strains (comprising multiple distantly related lineages including representatives of the well-known globally distributed ST14, ST23, ST86 and ST258). KPN_04246 encodes a protein that catalyses two reactions, ACODA, acetylornithine deacetylase, and NACODA, N-acetylornithine deacetylase, both of which may also be encoded by the product of KPN_01464 (homologs of this gene were identified in only those genomes that were not associated with loss of growth phenotype). Comparison of the *K. pneumoniae* strain CG43 (ST86) chromosome lacking KPN_01464 to *K. pneumoniae* MGH78578 harbouring KPN_01464 showed that the former contained a ∼10 kbp deletion resulting in the loss of KPN_01464. This deletion was replicated in the *K. africana* 200023T genome and the remaining 12 *K. pneumoniae* genomes that lacked KPN_01464 (≤33.24% identity for the best bi-directional BLASTp hit, no alternate genes encoding putative acetylornithine deacetylases/N-acetylornithine deacetylases were identified in KEGG).

Finally, we investigated the two gene deletions (KPN_02238 and KPN_00456) resulting in predicted loss of growth on all substrates in only *K. pneumoniae* NJST258-1. KPN_02238 encodes the protein responsible for catalysing PRPPS (phosphoribosylpyrophosphate synthetase), for which no redundant genes were included in any of our KpSC models. This reaction converts alpha-D-ribose 5-phosphate to 5-phospho-alpha-D-ribose 1-diphosphate, a key substrate utilised as input for 14 downstream reactions. While the *K. pneumoniae* MGH78578 reference model contains a redundant pathway to support this conversion, one of the required reactions (R15BPK, catalysed by a ribose-1,5-bisphosphokinase) was missing from the NJST258-1 model because the associated genome lacked a homolog of KPN_04492 (best bi-direction BLASTp hit 26.19% identity), whereas all other genomes contained a homolog of this gene. Further investigation showed that the NJST258-1 chromosome was missing a ∼17 kbp region compared to MGH78578. In the NJST258-1 chromosome, this region, which included KPN_04492, was replaced by the insertion sequence IS*1294* (99% nucleotide identity). Unfortunately, we were not able to identify a similar deficiency to explain the strain-specific loss of growth phenotype associated with KPN_00456, which encodes a protein implicated in 14 distinct reactions.

## Discussion

Here we present an updated GEM for *K. pneumoniae* MGH78578 plus novel GEMs for 36 members of the KpSC, capturing all seven taxa and representing the first reported GEMs for the *K. variicola* (subsp. *variicola* and *tropica*), *K. quasipneumoniae* (subsp. *quasipneumonaie* and *similipneumoniae*), *K. quasivariicola* and *K. africana* species. All models were validated and curated by comparison of predicted and true growth phenotypes, and had a median accuracy of 95.7% (range 88.3 - 96.8%), higher than estimated for the previously published *K. pneumoniae* MGH78578 (84%) and KPPR1 (79%) models.

Our *in silico* growth phenotype predictions for a diverse set of substrates highlighted variability among strains within the *K. pneumoniae* species, as has been indicated by previous smaller scale GEM comparisons and phenotypic comparisons (Norsigian et al. 2019a; Blin et al. 2017; Brisse et al. 2009; Henry et al. 2017). Similar variability was also indicated within and between the other species in the KpSC (**Fig. 3**). Carbon substrates were associated with the greatest diversity; a total of 145 substrates (53%) predicted to support growth of all 37 strains and 20 (7%) predicted to support growth of 1-36 strains each (**Fig. 2**). These predictions were consistent with the observed reaction variability, where the highest proportion of accessory reactions was identified among those associated with carbohydrate metabolism (16%, **Fig. 1**). This is consistent with a previous pan-genome analysis of 328 *K. pneumoniae* which indicated that ∼50% of the total gene-pool predicted to encode proteins with metabolic functions were specifically associated with carbohydrate metabolism (Holt et al. 2015). This trend is also consistent with previous studies of the closely related species, *Escherichia coli*, which demonstrated carbohydrate metabolism as the most diverse category for this organism (Fang et al. 2018; Monk et al. 2013).

The extent of diversity reported for *E. coli* and *Salmonella* spp. (Seif et al. 2018) was much higher than reported here for KpSC. We propose two likely explanations for these differences: i) the current analysis for KpSC comprises just 37 strains, compared to 55 and 110 strains included in the *E. coli* studies (Fang et al. 2018; Monk et al. 2013), and 410 in the *Salmonella* study (Seif et al. 2018). With greater sample size we expect to capture greater gene content diversity (Tettelin et al. 2008), including genes associated with metabolic functions that drive metabolic diversity (as was shown to be the case for *Salmonella* spp. (Seif et al. 2018)); ii) our draft KpSC strain-specific models were generated using the reference-based protocol (Norsigian et al. 2019b), where homology search is used to identify genes in the reference model that are absent from the strain of interest and are therefore removed from the strain-specific model. We added novel genes/reactions to the models based on comparison of predicted vs observed growth phenotypes and manual sequence/literature search, but we did not conduct an automated screen to identify additional genes that are present in the novel strain collection. The latter approach is expected to reveal further diversity, but it requires significant manual curation and validation to ensure the high-quality status of the models is maintained, and is therefore beyond the scope of the current study.

In addition to growth capabilities, our analyses revealed considerable variation in terms of predicted gene essentiality, as has been implicated for other bacterial species (Breton et al. 2015; Poulsen et al. 2019; Rousset et al. 2021; Tong et al. 2020). Specifically, our data indicate that i) deletion of a single core gene in a given strain may result in loss of growth on all, none or only a subset of growth substrates; and ii) the impact of such deletions may vary considerably between strains (**Fig. 4**). Amongst genes where deletion was predicted to have variable impact, most were associated with the loss of growth for only a small number of substrates in the impacted strains. However, four genes were associated with predicted loss of growth on ≥143 of 145 substrates for between one and 14 strains each. In two cases (genes KPN_03428 and KPN_04246), the impacted strains were missing redundant genes that were present in the MGH78578 reference model, i.e., those encoding proteins with the same functional annotation as the deleted gene. Comparisons of the chromosomes of these strains suggested that the genes were lost via large scale chromosomal deletions (5-10 kbp). One of these deletions was uniquely conserved among strains belonging to *K. quasipneumoniae* subsp. *quasipneumoniae*, suggesting that it may have occurred in the most recent common ancestor of this subspecies and has been inherited via vertical descent, with evidence from additional public genome data pointing towards recent re-acquisition of this region in some lineages. The other chromosomal deletion was found among a distantly related subset of *K. pneumoniae* as well as the single *K. africana* isolate, and therefore its distribution cannot be explained by simple vertical ancestry. Rather, we speculate that this deletion has been disseminated horizontally via chromosomal recombination, as is known to occur frequently among *K. pneumoniae* (Wyres et al. 2019; Bowers et al. 2015) and has been reported between KpSC species (Holt et al. 2015).

Deletion of two genes (KPN_02238 and KPN_00456) resulted in the loss of growth on all substrates for only a single strain (*K. pneumoniae* NJST258-1). This strain is of particular interest because it was associated with the highest number of deletion phenotypes (**Fig. 4**), and it belongs to ST258, a globally distributed cause of carbapenem-resistant *K. pneumoniae* infections (Wyres et al. 2020; Bowers et al. 2015). We were unable to identify the cause of this rare knockout phenotype (lacking adenylate kinase, encoded by KPN_00456), which converts D-ribose 1,5-bisphosphate to 5-phospho-alpha-D-ribose 1-diphosphate at the cost of 1 ATP. Comparison of the metabolic networks of NJST258-1 and MGH78578 indicated that NJST258-1 was lacking an additional reaction pathway (phosphoribosylpyrophosphate synthetase) present in MGH78578, allowing an alternative means of 5-phospho-alpha-D-ribose 1-diphosphate production in the absence of ribose-1,5-bisphosphokinase. Further investigation showed that the NJST258-1 chromosome was missing a ∼17 kbp region containing one of the genes required to express this redundant pathway, which had been replaced by an insertion sequence (IS). IS are frequently identified among *Klebsiella* and other Enterobacteriaceae where they are particularly associated with large plasmids and the dissemination of antimicrobial resistance (Che et al. 2021; Adams et al. 2016). The carbapenem-resistant *K. pneumoniae* lineage, ST258, has been associated with particularly high IS burden (Adams et al. 2016), and we hypothesise that such insertions contribute to the increased number of gene deletion phenotypes predicted for NJST258-1 compared to other *K. pneumoniae* strains. Further analyses will be required to assess whether this reduced redundancy is a general feature of ST258 or a trait specific to NJST258-1.

These findings indicate that KpSC can differ substantially in terms of metabolic redundancy. While we cannot exclude the possibility that the predicted knockout phenotypes might be rescued by products of non-orthologous genes that are not currently captured in our models, we note that at least for the examples described above, search of the KEGG database did not indicate any additional known redundant metabolic pathways. Additionally, our findings are consistent with a recent experimental exploration of gene essentiality in *E. coli* (Rousset et al. 2021), which showed that 7-9% of ∼3,400 conserved genes were variably essential among 18 *E. coli* strains grown in three different conditions. Importantly, genomic comparisons of these *E. coli* implicated a key role for horizontal gene transfer in driving strain-specific essentiality patterns and redundancies through the mobilisation of homologous or analogous genes and/or those driving epistatic interactions (Rousset et al. 2021).

Taken together our findings highlight the importance of strain-specific genomic variation in determining strain-specific metabolic traits and redundancy. More broadly, these analyses demonstrate the value of an organism investing in redundant systems, either through i) encoding multiple genes capable of performing the same reaction, or through ii) encoding multiple, alternative pathways for producing key metabolites from different substrates. Given what is known about the extent of genomic diversity among *K. pneumoniae* and the broader KpSC (Holt et al. 2015; Wyres et al. 2019; Thorpe et al. 2021), it is clear that studies seeking to understand the metabolism of these species – e.g., for novel drug design, or to identify novel virulence and drug resistance determinants – should include a diverse set of strains. In this regard, we anticipate that the GEMs, growth predictions and single gene deletion predictions presented here will provide a valuable resource to the *Klebsiella* research community. As exemplified for the *E. coli* K-12 reference strain, such resources can be continually improved and expanded to maximise their utility and facilitate biological discovery for years to come (Schilling et al. 1999; Monk et al. 2017).

## Methods

### Genome collection

The 37 strains used in this study were sourced from two previous studies (Blin et al. 2017; Rodrigues et al. 2019). Eight strains had completed genome sequences already publicly available, generated using various sequencing and assembly methods (see **Supplemental Table 1** for details). For the remaining 29 strains, short- and long-read sequencing was conducted as follows. Genomic DNA was extracted from overnight cultures, using GenFind v3 reagents (Beckman Coulter). The same DNA extraction was used for both Illumina and MinION libraries. Illumina sequencing libraries were made with Illumina DNA Prep reagents (20018705) and the Illumina Nextera DNA UD Indexes (20027217) as per manufacturer’s instructions with one major deviation from described protocol; reactions were scaled down to 25% of recommended usage. Illumina libraries were sequenced on the NovaSeq platform using the 6000 SP Reagent Kit (300 cycles; 20027465). Long-read sequencing libraries were prepared using the ligation library kit (LSK-109, ONT) with native barcoding expansion pack (EXP-NBD104 and NBD114, ONT). The library was run on a R9.4.1 MinION flow cell, and was base called with Guppy v3.3.3 using the dna_r9.4.1_450bps_hac (high-accuracy) basecalling model.

The Illumina and MinION read data were combined to generate completed genomes for n=28/29 strains with Unicycler v0.4.8 (Wick et al. 2017) using default parameters. SB610 could not be assembled into a completed genome using this approach, so we used Trycycler v0.3.3 (Wick et al. 2021) to combine 12 independent long-read only assemblies into a single consensus assembly. The 12 assemblies were generated from 12 independent subsets of the long reads (randomly selected) at 50x depth, which were assembled with one of three assemblers (n=4 assemblies each): Flye v2.7 (Kolmogorov et al. 2019), Raven v1.1.10 (Vaser and Šikić 2021) and Miniasm v0.3 (Li 2016). The final consensus assembly was then polished with the long reads using Medaka v1.1.3 (https://github.com/nanoporetech/medaka) followed by three rounds of polishing using the Illumina reads with Pilon v1.23 (Walker et al. 2014). All 37 completed genomes were annotated with Prokka v1.13.3 (Seemann 2014), using a trained annotation model (created using 10 genomes with Prodigal v2.6.3 (Hyatt et al. 2010)). All genomes were analysed with Kleborate v2.0.3 (Lam et al. 2021) to obtain ST and other genomic information (see **Supplemental Table 1**).

### Phenotypic testing

We utilised the Biolog growth phenotypes for 190 carbon substrates generated previously (Blin et al. 2017; Rodrigues et al. 2019). As determined in Blin et al., a maximum value in the respiration curve of ≥150 was used to indicate growth, whilst a value of <150 indicated no growth. We performed additional phenotypic tests on six carbon substrates; two which were not available on Biolog, 3-(3-hydroxy-phenyl)propionate (Sigma Cat Number PH011597) and 3-hydroxycinnamic acid (CAS Number 14755-02-3); and four Biolog substrates for which we required further evidence, gamma-amino butyric acid (CAS Number 56-12-2), L-sorbose (CAS Number 87-79-6), D-galactarate (CAS Number 526-99-8), and tricarballylate (CAS Number 99-14-9).

Overnight cultures of all 37 isolates were grown in M9 minimal media (2x M9, Minimal Salts (Sigma), 2 mM MgSO4 and 0.1 mM CaCl2) plus 20 mM D-glucose, at 37°C, shaking at 200 RPM. Each carbon source substrate solution was prepared to a final concentration of 20 mM in M9 minimal media, pH 7.0. Then, 200 μL of each substrate solution was added to separate 96-well cell culture plates (Corning) and 5 μL of overnight cultures added to the wells, diluted to McFarland standard of 0.4 – 0.55. Negative controls were included on every independent plate and included i) no substrate solution controls (20 mM M9 minimal media) and ii) no isolate controls but 20 mM substrate solution. For positive controls, each isolate was also grown independently in M9 minimal media containing 20 mM D-glucose. Every growth condition was performed in technical triplicate. Plates were then sealed with AeraSeal film (Sigma), then grown aerobically for 18 hours at 37°C, shaking at 200 RPM. Plates were then read using the FLUOstar Omega plate reader (BMG Labtech) using Read Control version 5.50 R4, firmware version 1.50, using 595 nm absorbance after 30 seconds of shaking at 200 RPM. No isolate controls were used as blanks for to generate the OD value for each technical replicate, then mean calculated to obtain the OD value. To determine growth/no growth using the OD method, we calculated the mean OD for growth on a particular substrate for each strain at 24h, and subtracted from this the OD value of M9 media alone. Subsequently, for each carbon substrate we divided the mean OD value for a strain by the mean OD for that strain in M9 media alone to get an OD fold change. OD fold changes ≥2 were considered sufficient evidence of growth (**Supplemental Figure 1**).

### Creating and curating strain-specific GEMs

Using the method outlined by Norsigian et al (Norsigian et al. 2019b), we extracted and translated all CDS from each genome and used bi-directional BLASTp hits (BBH) to determine orthologous genes compared to the reference *K. pneumoniae* MGH78578 GEM (iYL1288) (Liao et al. 2011). Genes with at least 75% amino acid identity were considered orthologous. Genes and their reactions that did not meet this threshold were removed from their respective models.

During GEM creation, we discovered that the original biomass function (BIOMASS_) in iYL1288 required the production of both rhamnose, which is a component of the capsule in *K. pneumoniae* MGH 78578, as well as UDP-galacturonate and UDP-galactose, which are components of the variable O antigen. As both the capsule and O antigens are known to differ greatly between strains (Wyres et al. 2016; Follador et al. 2016), we created a new biomass function (BIOMASS_Core_Oct2019) that no longer required the associated metabolites dtdprmn_c, udpgalur_c and udpgal_c.

To validate each GEM against its respective phenotypic growth results, we used flux based analysis (FBA) implemented in the COBRApy framework (Ebrahim et al. 2013) to simulate growth of each GEM in M9 media with all possible sole carbon, nitrogen, phosphorous or sulfur substrates. The updated BIOMASS function, BIOMASS_Core_Oct2019, was used as the objective to be optimised. M9 media was defined by setting the lower bound of the cob(I)alamin exchange reaction to -0.01, and the lower bound of the following exchange reactions to -1000: Ca2+, Cl-, CO2, Co2+, Cu2+, Fe2+, Fe3+, H+, H20, K+, Mg2+, Mn2+, MoO4 2-, Na+, Ni2+, Zn2+. To predict growth on alternate carbon substrates, we set the lower bound of glucose to zero (to prevent the model utilising this as a carbon source), and then set the lower bound of all potential carbon substrates to -1000 in turn. The carbon substrate was considered growth supporting if the predicted growth rate was ≥0.001.

While identifying carbon substrates, the default nitrogen, phosphorous and sulfur substrates were ammonium (NH4), inorganic phosphate (HPO4) and inorganic sulfate (SO4). Prediction of nitrogen, phosphorus and sulfur supporting substrates was performed in the same way as carbon, but setting glucose as the default carbon substrate.

We matched predictions and phenotypic growth data for all strains for 94 distinct carbon substrates. These data were used to i) curate and update the models; and ii) estimate model accuracy. Where we had evidence of phenotypic growth but a lack of simulated growth, we attempted to identify the missing reactions using gene homology searches and literature searches in related bacteria (see **Supplemental Table 3** for a full list of reactions added and the evidence for each). During this process it became apparent that the directionality of the following transport reactions in the original iYL1288 GEM were set to export the compound from the cell, rather than allow uptake (TARTRtex, SUCCtex, FORtex, FUMtex, THRtex, ACMANAtex, MALDtex, ABUTtex, AKGtex). Each of these reactions were updated to be reversible (bound range -1000 to 1000), restoring the ability for the model to utilise the associated compounds.

Strain model accuracy was determined by calculating the percentage of true positive and negative compounds, as well as calculating Matthew’s correlation coefficient using the following formula (TP = true positive; TN = true negative; FP = false positive; FN = false negative):

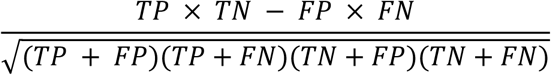

### Gene essentiality for growth on core carbon substrates

To determine which genes were essential for growth in each core carbon substrate (n=145) for each strain, we used the *single_gene_deletion* functions in COBRAPy (Ebrahim et al. 2013). For each GEM, on every core carbon substrate we simulated growth in M9 media with that substrate as the sole carbon source using FBA (as described above), but with one gene knocked out using the *single_gene_deletion* function. Each gene was knocked out in turn, and optimised biomass values ≥0.001 were considered positive for growth.

Four gene-substrate combinations were selected for further investigation by interrogation of the model gene-protein-reaction rules and search of the KEGG database (Kanehisa et al. 2002) using KofamKOALA (Aramaki et al. 2019) for redundant genes/pathways. Where relevant, pairwise chromosomal comparisons were performed using BLASTn (Camacho et al. 2009) and visualised using the Artemis Comparison Tool (Carver et al. 2005). The putative insertion sequence was identified by BLASTn search of the ISFinder database (Siguier et al. 2006).

### Core genome phylogeny

The core genome for the set of 37 genomes was determined using panaroo v1.1.2 (Tonkin-Hill et al. 2020) in strict mode with a gene homology cutoff of 90% identity, which generated a core gene alignment consisting of 3717 genes with 75,899 variable sites. We generated a phylogeny using this core gene alignment with IQTree v2 (Minh et al. 2020), which selected GTR+F+I+G4 as the best-fit substitution model. The resulting phylogeny was visualised using *ggtree* (Yu et al. 2017) in R.

## Supplemental Legends

**Supplemental Figure 1: Distribution of OD fold changes for growth on six carbon substrates**. Each panel is a substrate showing total number of strains (y axis) with a particular OD fold change (x axis). Red line indicates OD fold change of 2, fold changes greater than this value were considered sufficient evidence of growth.

**Supplemental Table 1: Strain collection in this study**. Genome information, model information.

**Supplemental Table 2: Growth/no growth for each strain-specific GEM in all possible substrates**. Prefix in front of the substrate indicates what source type it belongs to, C=carbon; N=nitrogen; P=phosphorous; S=sulfur.

**Supplemental Table 3: Details of reactions added to each strain-specific GEM**. Evidence level is in four categories (1-4) as per (Thiele and Palsson 2010): 1, no evidence available, reaction required for modelling; 2, evidence for gene function (genome annotation or SEED annotation), or indirect evidence based on phenotypic data; 3, direct and indirect evidence for gene function such as knockouts or expression analysis; 4, direct evidence of gene function and biochemical reaction.

**Supplemental Table 4: Details of model predictions vs phenotypic growth for all 94 substrates in each strain-specific GEM**.

**Supplemental Table 5: Growth phenotype predictions for all possible single-gene deletions in all strains, for growth on 145 carbon substrates**.

**Supplemental Table 6: Genes predicted to be essential for growth on all core carbon substrates for all strains**.

## Data Access

All completed genomes generated in this study have been deposited in GenBank under BioProject PRJNA768294 (accessions for individual genomes listed in **Supplemental Table 1**). All strain metabolic models generated in this study have been deposited in json format, along with the gene annotations used in the models, in figshare doi:10.26180/16702840.

## Acknowledgments

We thank Virginie Passet (Institut Pasteur) for assistance with Nanopore sequencing and Biolog data generation.

## Author Contributions

JH, KEH, JMM and KLW conceived the study and designed analyses. SB, CR, SLF and JSFL provided bacterial isolates and Biolog phenotype data. BV, LMJ, TH and CR generated novel sequence and/or phenotype data. JH, BV and KLW performed data analyses. JH, JMM, SB, KEH and KLW obtained funding. JH and KLW wrote the manuscript. All authors read, commented on and approved the manuscript.

