## Supplementary figures and images for "A curated collection of *Klebsiella* metabolic models reveals variable substrate usage and gene essentiality"

### Supplemental Figure 1

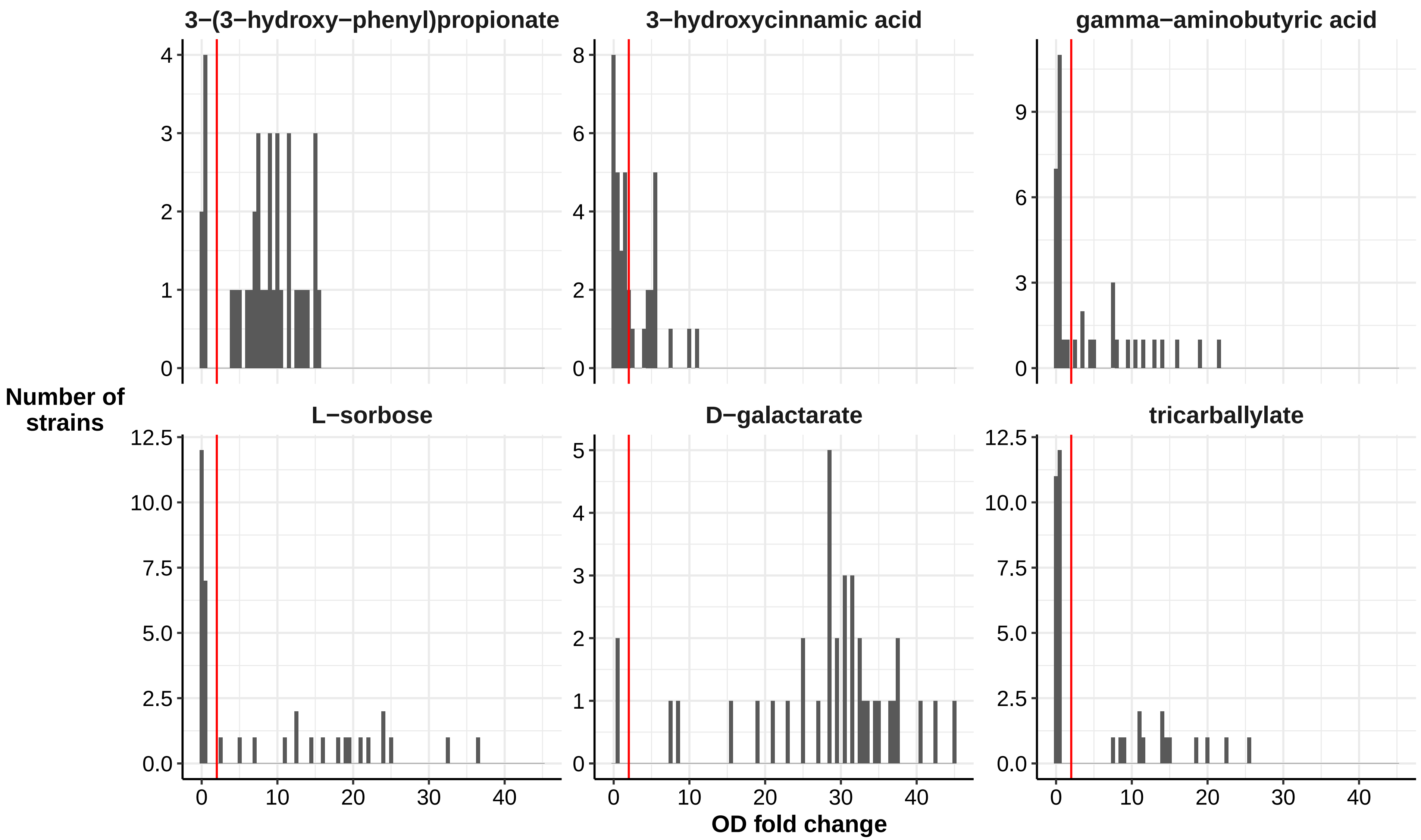
